# Transdiagnostic Neurobiological Biotypes of Trauma Timing: Data-driven approach of Childhood and Adulthood-Onset Trauma

**DOI:** 10.64898/2026.04.22.720234

**Authors:** Shilat Haim-Nachum, Chen Zhang, Kangyi Peng, Yuval Neria, Sigal Zilcha-Mano, Xi Zhu

**Affiliations:** School of Social Work, Tel Aviv University, Tel Aviv, Israel; Sagol School of Neuroscience, Tel Aviv University, Tel Aviv, Israel; Columbia University Department of Psychiatry & New York State Psychiatric Institute, New York, NY, USA; Department of Psychology, University of Haifa, Mount Carmel, Haifa, Israel; Department of Bioengineering, University of Texas at Arlington, Texas, USA

**Keywords:** Childhood trauma, adulthood trauma, rsFC, neurobiological markers, clusters

## Abstract

**Background:** The developmental timing of trauma exposure may critically shape neurobiological outcomes, yet distinctions between childhood-onset trauma (CT) and adulthood-onset trauma (AT) remain poorly understood.

**Aim:** This study explores whether trauma onset timing is associated with distinct resting-state functional connectivity (rsFC) pattern using data-driven approach.

**Methods:** Seventy-seven trauma-exposed individuals (*M*age=36.74 years) with post-traumatic stress disorder (PTSD), PTSD with major depressive disorder (MDD), and trauma-exposed healthy controls (TEHC) underwent resting-state fMRI. Of these participants, 15 with CT only, 17 with both CT and AT, and 47 with AT only. RsFC was calculated across the amygdala, hippocampus, nucleus accumbens (NAcc), the salience (SN), default mode (DMN), and frontoparietal networks (FPN). K-means clustering identified subgroups based on rsFC, with robustness assessed via bootstrapping, cross-validation, and replication using Gaussian Mixture Modeling. The identified clusters were compared on trauma timing, type, cumulative exposure, and clinical measures.

**Results:** A two-cluster solution provided the most stable fit. The two generated clusters were significantly different in CT-only prevalence (p < 0.05; Cramér’s V = 0.26, 95% CI). The CT cluster was marked by hyperconnectivity between amygdala–FPN, DMN–SN, NAcc–SN, and hippocampus–FPN relative to the AT cluster. Individuals with both CT and AT were evenly distributed across clusters. Clusters did not differ in PTSD or comorbid diagnoses, trauma type, or cumulative exposure.

**Conclusion:** Data-driven clustering revealed distinct neurobiological profiles differentiating CT and AT. CT was associated with hyperconnectivity across salience, reward, and regulatory circuits, supporting developmental timing as a determinant of brain network organization in trauma-exposed populations.

## Introduction

### Trauma Timing and Psychopathology

Trauma exposure is a recognized risk factor for multiple psychiatric and somatic conditions (1, 2, 3, 4). Growing evidence suggests that the *developmental timing* of trauma critically shapes long-term cognitive functions (5). A common distinction is between childhood-onset trauma (CT; <18 years) and adulthood-onset trauma (AT; ≥18 years). While this age demarcation is a simplification, it has grounding in developmental neuroscience: adversity during childhood and adolescence coincides with sensitive periods of brain maturation, stress-system calibration, and socioemotional development, rendering neural circuits particularly vulnerable to toxic stress (6, 7, 8, 9). Although developmental timing likely operates along a continuum rather than a fixed boundary, consistent with the “early-onset hypothesis,” CT has been repeatedly associated with enduring psychopathology, maladaptive functioning, and increased vulnerability to chronic psychiatric and health-related difficulties (10, 11, 12, 13, 14, 15). In contrast, AT has been most frequently studied in occupational or disaster contexts (e.g., military, first responders), with findings that remain heterogeneous, often depending on trauma type, context, and prior exposure (16, 17). These inconsistencies may reflect the influence of prior life stress and stress sensitization in adulthood, as well as differences in neural plasticity once brain systems are mature. Despite these differences, most neuroimaging studies have not separated CT from AT, limiting our ability to disentangle their distinct and independent neurobiological correlates.

### Neurobiological Heterogeneity and Trauma Timing

Resting-state functional connectivity (rsFC) provides a window into the intrinsic organization of large-scale brain networks following trauma. Altered coordination among systems supporting self-referential processing (default mode network; DMN), salience detection (salience network; SN), and cognitive control (frontoparietal network; FPN) has been repeatedly observed in trauma-exposed populations. However, findings remain inconsistent—CT has been linked to increased within-DMN coupling (e.g., PCC–vmPFC) and reduced amygdala–prefrontal connectivity (18, 19, 20), whereas AT studies more often associated with decreased DMN coherence and altered amygdala/hippocampal coupling with SN and FPN hubs (1, 21, 22). Such discrepancies likely reflect heterogeneity in sample composition, overlapping CT/AT histories, comorbidities, and analytic differences. Traditional case-control approaches/top-down designs (e.g., diagnosis-based group contrasts) may obscure effects related to developmental timing. Data-driven clustering methods offer a complementary approach by grouping individuals according to shared brain connectivity patterns rather than predefined diagnostic or exposure categories. While clustering approaches have begun to reveal neurobiological subtypes in trauma-related conditions (23, 24, 25), few studies have tested whether rsFC-based clusters correspond to CT versus AT histories independent of diagnosis or trauma load. Identifying neurobiological subtypes linked to developmental timing could improve mechanistic understanding, inform personalized interventions, and refine risk prediction models in trauma-exposed populations.

### Aim

In the present study we applied a data-driven clustering approach to examine whether trauma-exposed individuals cluster into subgroups characterized by distinct rsFC patterns, and whether these clusters reflect differences in the timing of trauma onset. We focused on three large-scale networks frequently implicated in trauma-related psychopathology: the DMN, SN, and ECN. We then examined whether the resulting clusters mapped onto the developmental timing of trauma exposure (childhood vs. adulthood) and whether they differed in clinical characteristics or trauma load. This approach aimed to reduce heterogeneity observed in prior work and to clarify whether trauma-onset timing is reflected in network-level brain connectivity. We hypothesized that data-driven clustering of rsFC would yield distinct subtypes, with CT-enriched subgroups characterized by hyperconnectivity of top-down control networks (e.g. FPN-amgydala, DMN-SN), and AT-enriched subgroups characterized by hypoconnectivity within these top-down control networks.

## Methods

### Participants

Seventy-seven of trauma exposed participants were recruited (age 18–60 years, *M*age = 36.74, SD = 12.38; 38% female) at the New York State Psychiatric Institute (NYSPI). The authors assert that all procedures contributing to this work comply with the ethical standards of the relevant national and institutional committees on human experimentation and with the Helsinki Declaration of 1975, as revised in 2013. All procedures involving human subjects/patients were approved by NYSPI Institutional Review Board (IRB), with participants providing written informed consent after receiving an explanation of the procedures. The IRB approval number is 7136. We characterized the type (interpersonal versus non-interpersonal trauma), severity levels, and timing of trauma exposure. Index trauma exposure met the Diagnostic and Statistical Manual of mental disorders, fifth edition (DSM-5) Criterion A of a traumatic event. Inclusion criteria were ability to give consent, English fluency, past experience of a traumatic event in childhood (age <18 years) and/or adulthood (age ≥18), and meeting the criterion A for PTSD of DSM-5. Trauma-exposed participants were excluded from the study if they met any of the following criteria: a prior or current diagnosis of schizophrenia, psychotic disorder, bipolar disorder, or dementia; A Hamilton Depression Rating (HAM-D) scale score greater than 25, indicating severe depression, depression-related impairment warranting pharmacotherapy, or combined medication and psychotherapy (26). Being at risk for suicide; A history of substance or alcohol dependence within the past six months, or abuse within the past two months; Any use of psychotropic medications, including antipsychotic, antidepressant, mood stabilizer, or stimulant medications, within the last four weeks prior to the study (six weeks for fluoxetine); Pregnancy; The presence of a medical illness that could interfere with assessment of response or biological measures, particularly during functional magnetic resonance imaging (fMRI); Having paramagnetic metallic implants or devices contraindicating magnetic resonance imaging, or any other non-removable paramagnetic metal in the body; Significant claustrophobia that impair participants’ ability to tolerate the MRI scanner environment.

### Instruments

Clinical evaluators administered the Structured Clinical Interview for DSM-5 Disorders (SCID-5; (27)) and the Clinician-Administered PTSD Scale for DSM-5 (CAPS-5; Weathers, Bovin (28)). The CAPS-5 is a 30-item structured interview assessing the frequency and intensity of each DSM-5 PTSD symptom on a 0–4 scale, yielding a total severity score (range 0–80). It demonstrates excellent interrater reliability and validity (29). Childhood-trauma questionnaire short form (CTQ-SF) was also administered; it is a 28-item screening inventory that assesses self-reported experiences of abuse and neglect in childhood and adolescence, it yields a total severity score (range-25 to 125)(30). Hamilton Anxiety Rating Scale (HAM-A) is a 14 items scale assessing anxiety symptom severity (range 0 – 56) (31).

Depressive symptoms were rated with the 17-item Hamilton Depression Rating Scale (HAM-D; (32)), a clinician-administered measure of depressive severity (range 0–52). Panic symptoms were evaluated with the Panic Disorder Severity Scale (PDSS; (33)), a 7-item self-report measure assessing the frequency, distress, and impairment associated with panic attacks (range 0–28). Lastly, the Temporal Experience of Pleasure Scale (TEPS) was administered. TEPS measures anticipatory and consummatory experiences of pleasures and it’s consisted of 18 items (range 18 to 108)(34). For clinical and demographic sample characteristics of the participant, see Table 1.

**Table 1:** Clinical and Demographic Information across CT-onset, AT-onset, and CT AT.

|  | Childhood-onset Trauma | Adulthood-onset Trauma | Both (CT and AT) | P-value (before correction) | P-value (FDR Corrected) |
| --- | --- | --- | --- | --- | --- |
| N | 15 | 45 | 17 |  |  |
| Female (%) | 7 | 12 | 10 | 4.79e-02 | 0.57 |
| Race |  |  |  | 3.92e-01 | 1 |
| African-American | 6 | 20 | 12 |  |  |
| Hispanics | 2 | 9 | 0 |  |  |
| Asian | 2 | 5 | 1 |  |  |
| Caucasian | 3 | 7 | 4 |  |  |
| Mix | 2 | 4 | 0 |  |  |
| CAPS | 22.53 | 17.4 | 20.47 | 0.55 | 0.67 |
| CTQ_TOT | 50.83 | 42.51 | 45.23 | 0.04 | 0.29 |
| HAM-A | 10.33 | 8.4 | 8.76 | 0.73 | 0.78 |
| HAM-D | 8.27 | 7.22 | 8.76 | 0.72 | 0.78 |
| PDSS_TOT | 8.87 | 4.19 | 4.41 | 0.03 | 0.29 |
| TEPS anticipatory | 38.4 | 40.72 | 43.06 | 0.43 | 0.62 |

CAPS, Clinician administered PTSD scale; CTQ, Childhood trauma questionnaire; HAM-A, Hamilton Anxiety Rating Scale-Anxiety; HAM-D, Hamilton Anxiety Rating Scale-Depression; PDSS, Panic Disorder Severity Scale; TEPS anticipatory, The Temporal Experience of Pleasure Scale-anticipatory

**Table 2:** Clinical and Demographic Information across two clusters.

|  | Cluster 1 (CT) | Cluster 2(AT) | P-value (before correction) | P-value (FDR corrected) |
| --- | --- | --- | --- | --- |
| N | 38 | 39 |  |  |
| Gender, N (%) |  |  | 0.93 | 1 |
| Male | 23(60.52%) | 25(64.10%) |  |  |
| Female | 15(39.47%) | 14(35.90%) |  |  |
| Race, N (%) |  |  | 0.64 | 1 |
| African-American | 18 (47.37%) | 20 (51.28%) |  |  |
| Hispanic | 7(18.42%) | 6 (15.38%) |  |  |
| Asian | 4(10.52%) | 1 (2.56%) |  |  |
| Caucasian | 8(21.05%) | 10 (25.64%) |  |  |
| Mixed | 1(2.63%) | 2 (5.13%) |  |  |
| Age, mean years (SD) | 38.4(13.0) | 35.1(11.7) | 0.243 | 0.378 |
| Education, mean years (SD) | 14.7 (2.37) | 14.6 (2.73) | 0.791 | 0.804 |
| Trauma type (CT only) | 12 | 3 | 0.03* | 0.48 |
| CAPS total severity scores | 31.58 | 16.46 | 0.18 | 0.39 |
| CTQ total | 49.23 | 45.32 | 0.38 | 0.49 |
| HAM-A | 10.87 | 6.90 | 0.05 | 0.39 |
| HAM-D | 9.10 | 6.46 | 0.12 | 0.39 |
| PDSS total | 6.08 | 4.33 | 0.22 | 0.39 |
| TEPS anticipatory total | 37.92 | 43.44 | 0.02* | 0.39 |

CAPS, Clinician administered PTSD scale; CTQ, Childhood trauma questionnaire; HAM-A, Hamilton Anxiety Rating Scale-Anxiety; HAM-D, Hamilton Anxiety Rating Scale-Depression; PDSS, Panic Disorder Severity Scale; TEPS anticipatory, The Temporal Experience of Pleasure Scale-anticipatory

### Neuroimaging Acquisition

A total of seventy-seven participants’ resting-state fMRI were collected using two scanners due to scanner upgrade. Fifty-three participants were scanned using a 3T General Electric MR750 scanner (GE Medical Systems, Waukesha, WI, USA) and 24 participants were scanned using a 3T GE PREMIER scanner (GE Medical Systems, Waukesha, WI, USA), equipped with a 32-channel receive-only head coil. For each participant, a high-resolution T1-weighted 3D BRAVO sequence was acquired using the following parameters: T1□=□450□mm, Flip angle□=□12°, field of view□=□25.6□cm, 256□×□256 matrix, slice thickness□=□1□mm. rs-fMRI images were acquired using T2*-weighted echo-planar images (EPIs) depicting the blood-oxygen-level dependent (BOLD) using the following parameters: TR=1.3 sec, TE=28 msec, FA =60°, FOV =19.2 cm, number of slices=27, slice thickness=4 mm. Participants were instructed to relax, remain awake, and lie still with their eyes open. Foam pads securing participants in the head coil were used to limit head movement during data acquisition.

### Neuroimaging Preprocessing

RsFC analyses were carried out using a seed-based approach implemented in the CONN-fMRI Functional Connectivity toolbox v13(35). ROI-to-ROI connectivity analysis was performed using 15 ROIs of SN, DMN, and ECN, amygdala, hippocampus and NAcc based on CONN default atlas, previously identified as important in childhood trauma and adulthood trauma. The mean BOLD time series was computed across all voxels within each ROI. Bivariate regression analyses were used to determine the linear association of the BOLD time series between each pair of regions for each individual. The resultant correlation coefficients were transformed into z scores using Fisher’s transformation to satisfy normality assumptions.

### Statistical Analyses

Multiple linear regressions were used to adjust all a priori selected 105 rsFC features for sex and type of scanner. To divide participants into homogeneous clusters according to rsFC, unsupervised machine learning algorithm k-means clustering method was used (36). K-means clustering seeks to identify clusters, using neuroimaging data, in which the total within-cluster variation is minimized. Each observation is assigned to a given cluster to minimize the sum of squared distances of the observation to their assigned cluster centers (37). We used the R function kmeans in package stat to implement the k-means clustering on the adjusted rsFC features.

To identify the optimal number of clusters that best fit the data, we used the average silhouette method to determine the optimal number of clusters that best fit the data (38). Average silhouette method computes the average silhouette, which quantifies how well an observation fits within its assigned cluster compared to other clusters, of observations for different tested values of k. The optimal number of clusters k maximizes the average silhouette compared to a range of possible values for k. To visualize the separation between the clusters, we conducted principal component analysis (PCA) and plotted the individual participants across the two principal components.

To characterize the clusters and identify the features that best differentiate between them, we conducted a two□sample t□test to compare the two clusters on ROI-to-ROI connectivity, demographic, and clinical characteristics.

To assess the robustness of the k-means solutions (for k = 2, 3, 4), the stability and consistency of the clusters assignment were assessed using the following methods: bootstrapping through examining how often a subject is repeatedly assigned into consistent cluster, simulation-based significance testing and permutation-based significance testing compute the null distribution of average silhouette (the average distance between each point to its own cluster and its nearest neighboring cluster), thereby assessing the strength of observed clustering compared to chance-level. Split-half reliability was used by splitting the participants into two equal sets, and applying clustering to each half, followed by adjusted rand index to assess the congruency across the generated clusters in each half. 10-fold cross-validation was used to partition the data, within each fold, 90% of the data was used to train for k-means clustering, then followed by assignment of remaining 10% to each cluster based on the trained dataset. Adjusted rand index was computed to be compared to the full-sample k-means solutions. To assess the robustness of k-means clustering solution, the analysis was repeated using Gaussian Mixture Modeling (GMM). GMM assumes that data exists in Gaussian distribution and assigns participants to cluster based on the maximum posterior probability of membership. GMM was applied to the same set of neuroimaging rsFC of the participants, with the number of clusters specified by k=2. The resulting participant assignment to respective clusters in GMM was then examined for their overlap with k-means generated clusters.

## Results

### Demographics and Clinical Characteristics

Demographic and clinical characteristics, including diagnoses, across the CT, AT, and both CT AT participants are presented in Table 1. The three groups did not differ in age, sex, race, and across clinical scales (Table 1). We also compared trauma exposure characteristics across the data-driven clusters. The number of interpersonal trauma events did not differ significantly between Cluster 1 (n = 23) and Cluster 2 (n = 16) (p = 0.16). Similarly, cumulative trauma exposure did not differ between clusters. Participants were divided into trauma-severity groups—mild (1–4 events), moderate (5–8), severe (9–12), and very severe (13–16) (for a similar dose-response approach, see Flinn, Hefferman-Clarke (39)—and the proportions across these categories were comparable between clusters (p = 0.412).

### Cluster Identification

The average silhouette method suggested that two clusters best fit the data (Supplementary Figure 1). Two clusters were identified using the k□means clustering method: Cluster 1 (N = 39) and Cluster 2 (N = 38). The separation of the two clusters is visualized in Supplementary Figure 4 in principal component space. The clusters are characterized by 15/39 PTSD in cluster 1 and 19/38 PTSD in cluster 2. By contrast, the two clusters significantly differed in the proportion of participants that experienced CT only, with cluster 1 having a higher proportion of CT (32%, 12/38), and cluster 2 having lower proportion of CT (8%, 3/39) and a higher proportion of AT-only individuals (67%, 26/39). There were no significant differences between two clusters in their proportions of participants with AT only or AT/CT comorbidity (Figure 1a).

**Figure 1a:**
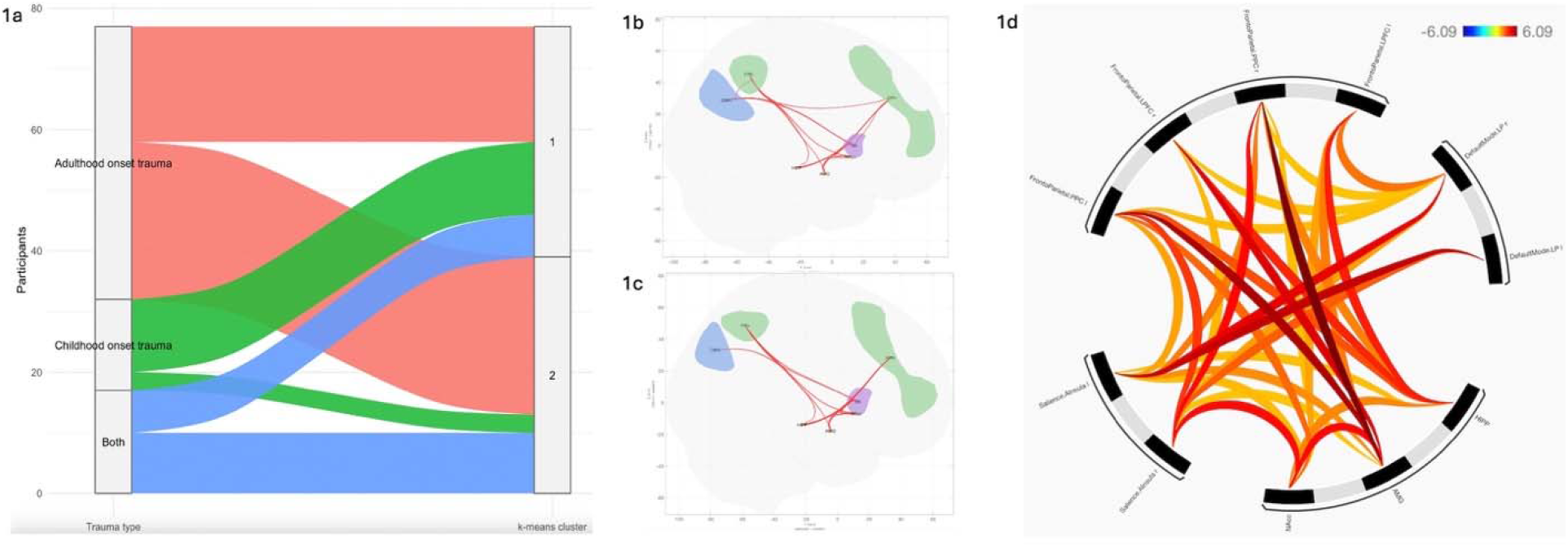
shows the distribution of CT, AT, both CT+AT participants across the two clusters, cluster 1 on top, labelled as 1, cluster 2 on the bottom, labelled as 2. 5b Cluster 1 ROI-to-ROI connectivity, 5c Cluster 2 ROI-to-ROI connectivity. 5c illustrate the connectivity differences between clusters The nodes are color-coded by functional network: SN (purple); limbic regions, including hippocampus (HIP), amygdala (AMG, middle), and nucleus accumbens (NAcc) (all in orange); DMN (blue); and FPN (green). Red edges indicate connections with significantly greater functional connectivity in Cluster 1 compared to Cluster 2, controlling for sex, age, and scanner. Displayed results correspond to clusters significant at *p*FDR < .05. 5c shows a circular connectogram of between-cluster differences in rsFC.

Out of 77 trauma-exposed participants, 18 met criteria for current PTSD diagnosis, 1 met criteria for lifetime PTSD diagnosis, 11 for current Major Depressive Disorder (MDD) and 18 for lifetime MDD, 11 for current Persistent Depressive Disorder (PDD) and 2 for lifetime PDD, 11 for current panic disorder (PD) and 2 for lifetime PD, 6 for SAD and 0 for lifetime SAD, 1 for current Obsessive Compulsive Disorder (OCD) and 1 for lifetime OCD, lastly, 0 for current Substance Use Disorder (SUD) and 7 for lifetime SUD. For more information on comorbid diagnosis of participants, see supplementary Table 1.

Two sample t□test analyses suggested that the two clusters did not differ in the proportion of PTSD, MDD, PDD, GAD, SAD, PD, OCD, and SUD (Supplementary Table 2), for both lifetime and current diagnosis. This suggests that data-driven clusters fail to identify diagnostic markers.

To ensure the cluster assignment was not confounded by the trauma type or severity, we conducted a group comparison of the interpersonal trauma and the cumulative trauma counts of participants within each cluster. The number of interpersonal trauma and the cumulative trauma counts were not significantly different across clusters.

### Cluster Brain Characteristics

The clusters differed significantly in the rsFC of amygdala-FPN, DMN-SN, Amygdala-NAcc, NAcc-SN, hippocampus-FPN, and SN-FPN (Figure 1c). Cluster 1 (CT) is characterized by hyperconnectivity within amygdala-FPN, DMN-SN, Amygdala-NAcc, NAcc-SN, hippocampus-FPN, and SN-FPN compared to cluster 2 (Figure 1b). However, cluster 1 (CT) is characterized by lower ability to anticipate pleasure/greater anhedonia (TEPS_AntTOT), though non-surviving of FDR Corrections (p < 0.05, p.adj = 0.388).

### Cluster Stability and Consistency

To decide on optimal k and cluster stability, we used bootstrapping, elbow, silhouette, adjusted rand index. Using bootstrapping, the average stability index for k = 2 is ∼0.58 for the two clusters, while the average stability index for k = 3 is approximately 0.72 across the three clusters. Average silhouette width for k = 2 (∼0.06) and k=3 (∼0.06) were not significantly different (Supplementary figure 2), whereas average silhouette width for k = 4 was lower (∼0.05) (Supplementary Figure 1). Despite the k = 3 being the highest average silhouette score, the permutation-based significance testing further confirmed the weak silhouette indices, permutation p-values demonstrate that the separation was not above chance (p = 0) (Supplementary Figure 3). Mean adjusted rand index using split-half reliability for k = 2 (∼0.07) was less stable than k = 3 (∼0.33), and mean adjusted rand index using 10-fold CV for k = 3 (0.95) was higher than k = 2 (0.66); this suggests that held out points were consistently assigned to the same clusters within the three clusters solution.

### Sensitivity Analysis

To examine the robustness of the k-means clustering solution, the analysis was repeated with the Gaussian mixture model (GMM) and similar findings were revealed. The GMM yielded 2 clusters. To assess the similarity of the 2 clusters generated using GMM and k-means, the groupings of patients’ data derived separately using k-means clustering and GMM were compared to each other for the overlap of participant assignment. We found a 100% overlap between cluster 1 in GMM and 100% of cluster 2 in k-means (n = 38), similarly, a 100% overlap of cluster 2 in GMM and cluster 1 of k-means (n= 39). The two clusters generated using GMM also demonstrated a significantly different proportion of CT, characterized by 12 CT in cluster 1 and 3 CT in cluster 2, which is congruent with the results from k-means clustering. Chi-square analysis was repeated for GMM-generated clusters, reporting significant differences (p < 0.05).

## Discussion

This study is among the first to apply a data-driven clustering approach to examine whether rsFC patterns distinguish individuals with childhood-onset trauma (CT) from those with adulthood-onset trauma (AT). Across two clustering methods, we identified two rsFC-defined subgroups that differed in the proportion of CT and AT histories and were characterized by distinct network-level connectivity profiles. The convergence of findings across analytic approaches suggests that developmental timing of trauma may leave a measurable imprint on large-scale functional brain organization, even when diagnostic categories are held constant.

The subgroup enriched for CT was characterized by hyperconnectivity among salience, control, and reward networks, including amygdala–FPN, DMN–SN, hippocampus–FPN, amygdala–NAcc, and SN–NAcc (Figure 2). This pattern aligns with prior evidence linking early adversity to atypical maturation of large-scale networks and heightened top-down demands on emotion regulation (6, 18, 19). Such hyper-integration may reflect chronic allostatic load and inefficient regulation of threat and reward signals. In mechanistic terms, early trauma occurs during periods of rapid neurodevelopment when salience–control–reward networks are still maturing, which may bias these circuits toward hypervigilance and sustained emotional engagement. This pattern could represent an adaptive but costly calibration of neural systems toward over-detection of threat and dysregulated reward valuation (40). At the same time, heterogeneity in prior CT findings—ranging from hyper-to hypoconnectivity—underscores the complexity of developmental timing effects and the likely contributions of trauma severity, comorbidity, and sample characteristics (7). By identifying a CT-enriched profile through an unsupervised, data-driven approach, our findings suggest that timing of exposure may shape connectivity patterns in ways not captured by traditional diagnostic categories.

**Figure 2:**
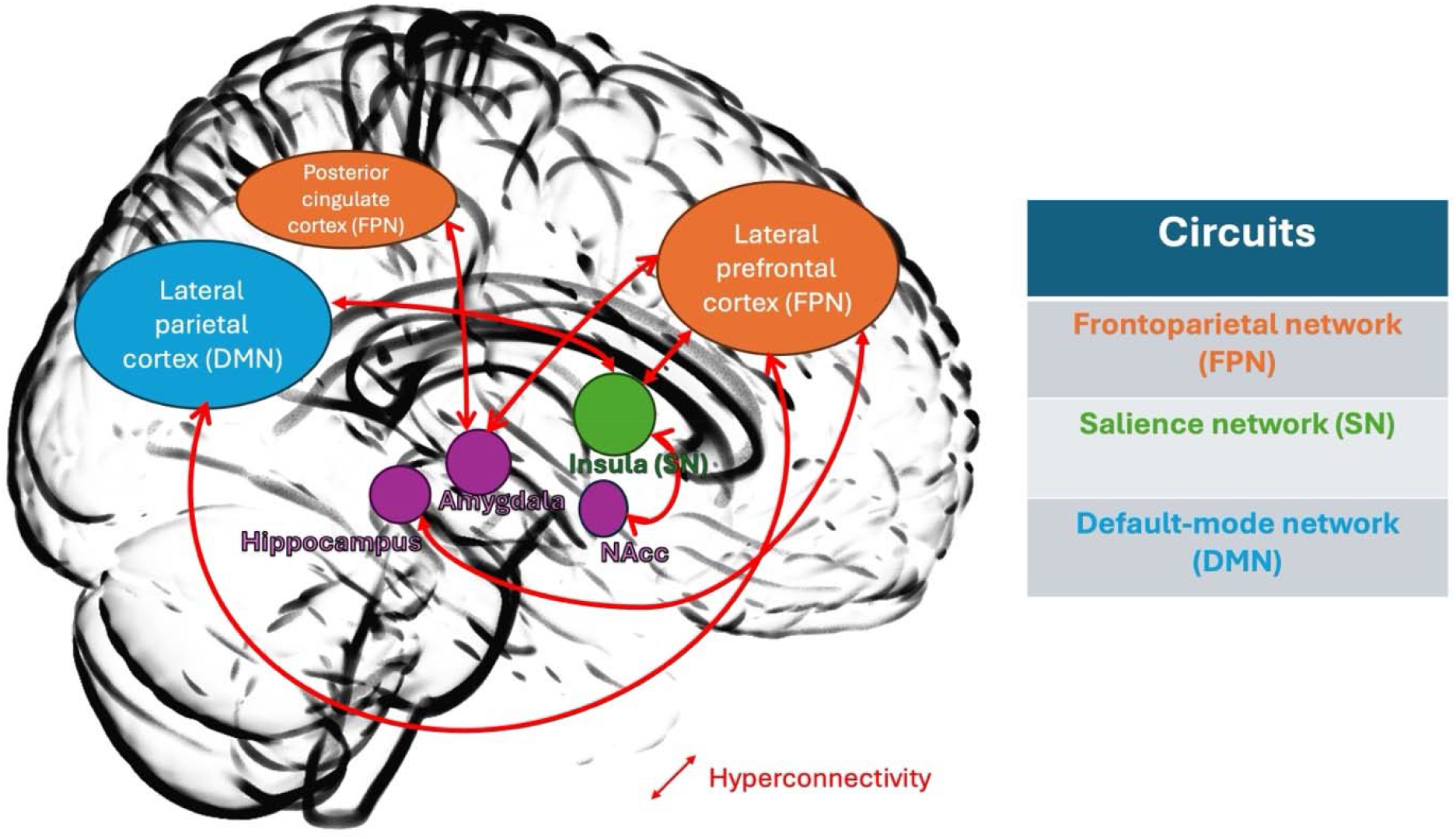
This is a visualization of rsFC hyperconnectivity within the CT cluster. Hubs within FPN are in orange, SN hub is in green, and DMN hub is in blue. Subcortical regions such as hippocampus, Amygdala, and NAcc are in purple. Hyperconnectivity observed in the following regions: lateral prefrontal cortex of FPN to lateral parietal cortex of DMN, lateral parietal cortex (DMN) to insula (SN), posterior cingulate cortex (FPN) to amygdala, lateral prefrontal cortex (FPN) to amygdala, hippocampus to lateral prefrontal cortex (FPN), insula (SN) to lateral prefrontal cortex (FPN), and insula (SN) to Nucleus Accumbens (NAcc). Bolded red arrows reflect the hyperconnectivity between regions.

In contrast, the AT-enriched cluster showed relative hypoconnectivity across the same circuits. This pattern aligns with reports of reduced DMN integrity and weakened coupling between limbic and control networks in adult-onset trauma samples (21, 22). Functionally, trauma encountered after neural systems have largely matured may induce compensatory disengagement or network decoupling, reflecting diminished integration of emotional salience and executive control processes. Taken together, these bidirectional findings suggest that CT and AT may represent distinct mechanistic pathways to trauma-related psychopathology: early trauma may sensitize salience and control systems to overreact to threat and social cues, whereas later trauma may impair network integration and flexibility. These pathway differences may carry implications for treatment—targeting hyperarousal and reward recalibration in CT-related presentations and strengthening engagement and cognitive integration in AT-related ones.

Exploratory analyses suggested that the CT-enriched cluster was associated with lower anticipatory pleasure compared to the AT-enriched cluster, although this effect did not survive correction for multiple comparisons. Nevertheless, the pattern is consistent with the observed fronto-striatal hyperconnectivity and prior links between early adversity, altered reward circuitry, and anhedonia (40, 41). While preliminary, this convergence highlights the potential utility of integrating dimensional reward-related traits into clustering frameworks to better capture trauma-related subtypes.

Two distinct clustering methods (k-means and GMM) on the neuroimaging data yielded clusters that overlapped 100%, supporting a stable differentiation between CT and AT groups rather than algorithm-specific noise. However, the low silhouette width and modest adjusted Rand index values indicate weak categorical separation, suggesting caution in inferring discrete subtypes: CT- and AT-related profiles may reflect partially overlapping network configurations along a broader trauma-related continuum. This dimensional interpretation complements, rather than contradicts, the categorical findings—it acknowledges that developmental timing exerts reliable but graded influences on neural organization. From this perspective, clustering helps identify anchor points along this gradient (CT-enriched vs. AT-enriched connectivity) while recognizing the inherent fluidity of trauma-related neural adaptations.

Several limitations should be noted. Subgroup sizes, particularly for CT, were small, limiting statistical power and stability. Data were collected on two scanners, and although scanner effects were adjusted for, residual confounding cannot be excluded. The cross-sectional design precludes causal inferences about whether the observed connectivity patterns reflect preexisting vulnerabilities, consequences of trauma exposure, or adaptive changes over time. Because we examined only static rsFC, our findings capture relatively stable network relationships but not their temporal dynamics. Future studies could extend this work by examining dynamic FC, regional homogeneity, or other derived resting-state metrics to capture additional aspects of local and time-varying network function. Finally, clinical and reward-related differences between clusters were modest and thus require replication in larger, independent samples.

Taken together, the present study provides preliminary evidence that developmental timing of trauma is reflected in distinct rsFC patterns. CT was linked to hyperconnectivity across salience, reward, and control circuits, whereas AT was associated with relative hypoconnectivity. These findings suggest that CT and AT may represent different mechanistic pathways to trauma-related psychopathology, with potential implications for tailoring interventions. At the same time, the findings highlight the potential utility of trauma timing as a transdiagnostic organizing dimension for understanding variability in brain function among trauma-exposed individuals.

Rather than mapping onto any single clinical diagnosis, the observed connectivity patterns may reflect underlying mechanisms that cut across diagnostic categories, offering insight into dimensions of vulnerability, resilience, and psychological functioning more broadly. This approach aligns with emerging frameworks that prioritize functional biomarkers and dimensional models of psychopathology over traditional categorical systems. Future research should replicate these findings in larger, longitudinal cohorts, integrate multimodal measures (e.g., structural connectivity, physiological indices, reward phenotypes), and examine whether CT-versus AT-related connectivity profiles predict differences in clinical course or treatment response.

## Acknowledgement

None

## Author Contribution

SHN and CZ, interpreted the data, drafted, and reviewed the manuscript

CZ and XZ conducted statistical analysis

CZ and KP designed and generated figures

YN, SZM, and XZ reviewed the manuscript

YN collected and acquired the data

