## Supplemental material for "Transdiagnostic Neurobiological Biotypes of Trauma Timing: Data-driven approach of Childhood and Adulthood-Onset Trauma"

  

**SUPPLEMENTARY INFORMATION**

Transdiagnostic Neurobiological Biotypes of Trauma Timing: Cluster-Based Differentiation of Childhood and Adulthood-Onset Trauma

**TABLE OF CONTENTS**

FIGURES ………………………………...……….…..….…..….…..….…..…..…..….  2-3

TABLES  ………………………………...……….…..….…..….…..….…..…..…..….   4-9

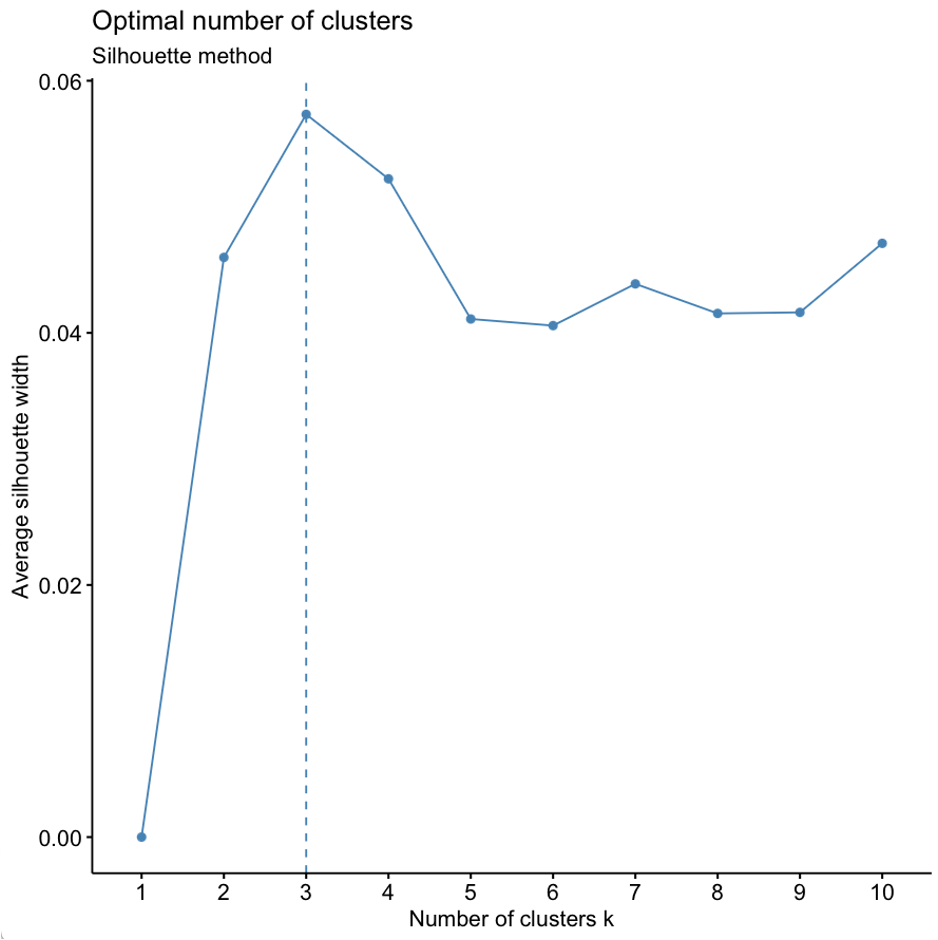

Figure 1. average silhouette width is plotted against the number of clusters. The highest silhouette value was observed at k = 3, suggesting three optimal clusters.

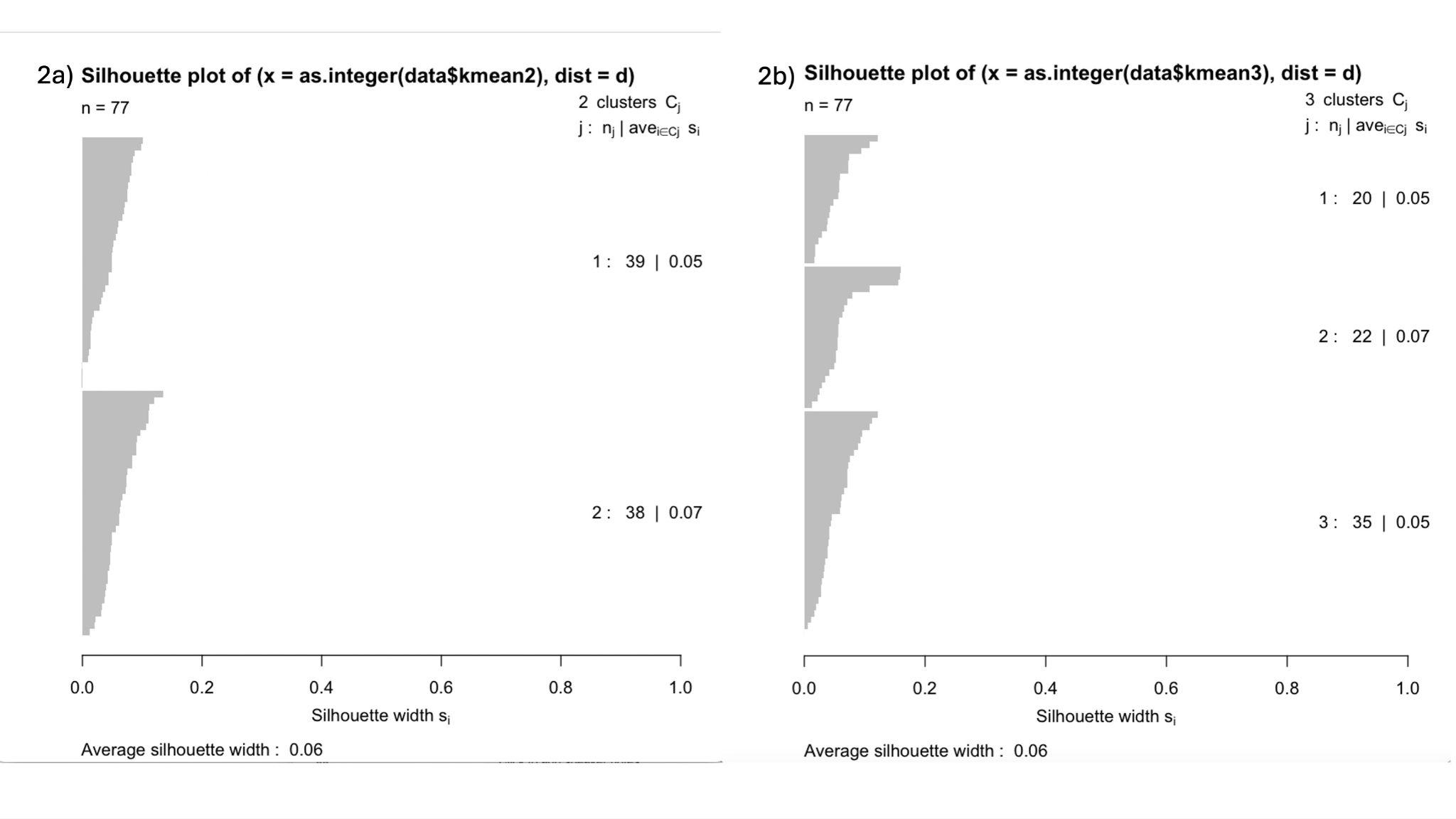

Figure 2. Silhouette plots illustrating clustering structure for 2-cluster k-means solution, and 3-cluster k-means solution. 2-cluster silhouette plots demonstrate that the average silhouette width for 2-cluster k-means solution (2a) and 3-cluster k-means solution is both 0.06 (2b).

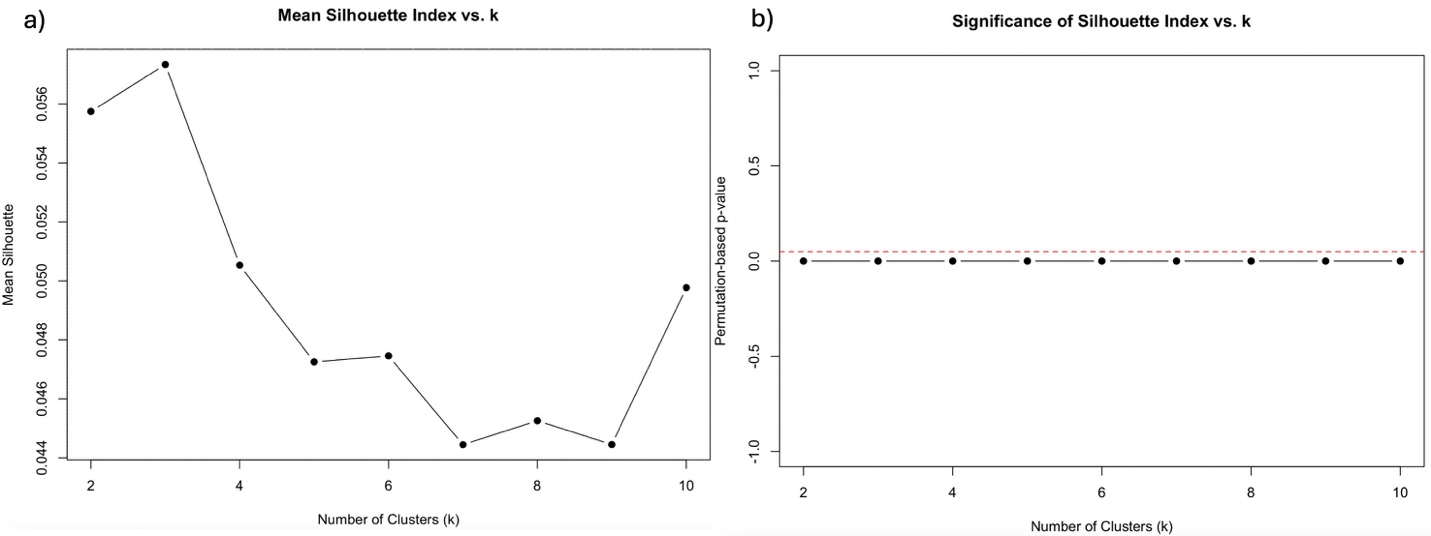

Figure 3. *Mean silhouette index and permutation-based significance of cluster solutions.*

**(a)** The mean silhouette index was calculated for k-means cluster solutions of 2 to 10. The average silhouette width peaked at 2-cluster k-means solution and 3-cluster k-means solution. The silhouette index declined starting from k = 4 to 10, indicating reduced clustering quality.

**(b)** The permutation-based p-values for each cluster solution revealed that the observed silhouette values were not significantly higher than those expected by chance (approximated zero –values for all k-cluster solutions), suggesting weak overall cluster separation in the data.

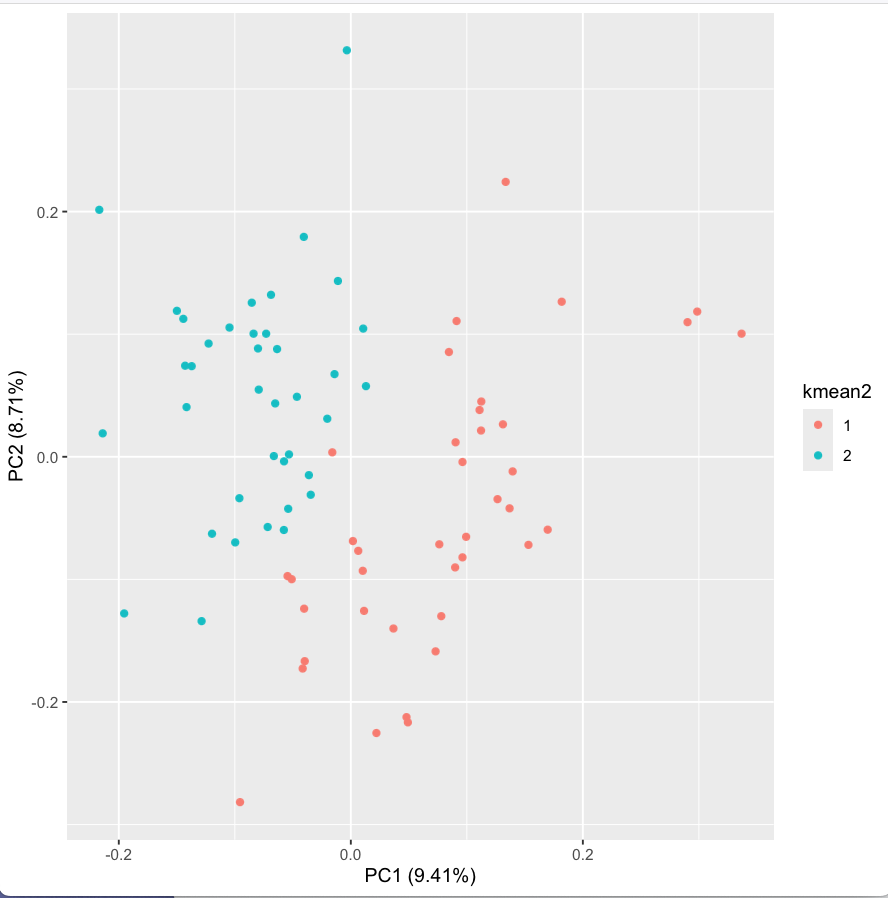
›

Figure 4 visualizes the separation of the two clusters, visualized in the space of first two principal components (PC1: 9.41%, PC2: 8.71%)

Table 1: Clinical and Demographic Information across CT-onset, AT-onset, and CT AT

|  | Childhood-onset Trauma | Adulthood-onset Trauma | Both (CT and AT) | P-value (before correction | P-value (FDR Corrected) |
| --- | --- | --- | --- | --- | --- |
| Sex |  |  |  | 4. 788685e-02 | 0.57 |
| Male | 8 | 33 | 7 |  |  |
| Female | 7 | 12 | 10 |  |  |
| MDD current count |  |  |  | 0.95 | 0.95 |
| Yes | 2 | 7 | 3 |  |  |
| No | 113 | 38 | 14 |  |  |
| MDD lifetime |  |  |  | 0.57 | 0.72 |
| Yes | 5 | 9 | 4 |  |  |
| No | 10 | 36 | 13 |  |  |
| PTSD current |  |  |  | 0.18 | 0.47 |
| Yes | 9 | 16 | 9 |  |  |
| No | 6 | 29 | 8 |  |  |
| PTSD lifetime |  |  |  | 0.17 | 0.47 |
| Yes | 0 | 0 | 1 |  |  |
| No | 15 | 45 | 16 |  |  |
| PDD current |  |  |  | 0.212 | 0.47 |
| Yes | 4 | 4 | 3 |  |  |
| No | 11 | 41 | 14 |  |  |
| PDD lifetime |  |  |  | 0.48 | 0.73 |
| Yes | 1 | 1 | 0 |  |  |
| No | 14 | 44 | 17 |  |  |
| GAD current |  |  |  | 1.58e-03 | 8.58e-03 |
| Yes | 3 | 0 | 0 |  |  |
| No | 12 | 45 | 17 |  |  |
| GAD lifetime |  |  |  | Not applicable | Not applicable |
| Yes | 0 | 0 | 0 |  |  |
| No | 15 | 45 | 17 |  |  |
| SAD current |  |  |  | 3.41e-01 | 0.63 |
| Yes | 2 | 4 | 0 |  |  |
| No | 13 | 41 | 17 |  |  |
| SAD lifetime |  |  |  | Not applicable | Not applicable |
| Yes | 0 | 0 | 0 |  |  |
| No | 15 | 45 | 17 |  |  |
| OCD Current |  |  |  | 0.70 | 0.82 |
| Yes | 0 | 0 | 1 |  |  |
| No | 15 | 45 | 16 |  |  |
| OCD Lifetime |  |  |  | 0.17 | 0.46 |
| Yes | 0 | 1 | 0 |  |  |
| No | 15 | 44 | 17 |  |  |
| Panic disorder current |  |  |  | 1.58e-04 | 1.7e-03 |
| Yes | 5 | 0 | 1 |  |  |
| No | 10 | 45 | 16 |  |  |
| Panic disorder lifetime |  |  |  | NA | NA |
| Yes | 0 | 0 | 0 |  |  |
| No | 15 | 45 | 17 |  |  |
| Substance abuse current |  |  |  | NA | NA |
| Yes | 0 | 0 | 0 |  |  |
| No | 15 | 45 | 17 |  |  |
| Substance abuse lifetime |  |  |  | 0.76 | 0.84 |
| Yes | 1 | 5 | 1 |  |  |
| No | 14 | 40 | 16 |  |  |
| ADHD current |  |  |  | 0.60 | 0.73 |
| Yes | 0 | 3 | 1 |  |  |
| No | 15 | 42 | 16 |  |  |
| ADHD lifetime |  |  |  | NA | NA |
| Yes | 0 | 0 | 0 |  |  |
| No | 15 | 45 | 17 |  |  |
| GAD-7_TOT | 9.2 | 5.51 | 5.82 | 0.13 | 0.35 |
| PCL_TOT | 33.07 | 22.02 | 25.59 | 0.2 | 0.45 |
| MASQ_TOT | 155.93 | 125.7 | 131.65 | 0.07 | 0.34 |
| SDSTOT | 15.73 | 9.12 | 10.18 | 0.08 | 0.34 |
| TEPS consummatory | 33.53 | 32 | 32.12 | 0.82 | 0.85 |

GAD-7, Generalized anxiety disorder-7; MASQ, Mood and Anxiety Symptom Questionnaire; SDS, Zung Self-Rating Depression Scale; TEPS consummatory, The Temporal Experience of Pleasure SCale-consummatory; SHAPS, Snaith-Hamilton Pleasure Scale

**Table 2: Clinical Information across two clusters**

|  | Cluster 1 (CT) | Cluster 2(AT) | P-value (before correction) | P-value (FDR corrected) |
| --- | --- | --- | --- | --- |
| Trauma type |  |  | 0.03 | 0.48 |
| Adult onset trauma only | 19 | 26 |  |  |
| Childhood onset trauma only | 12 | 3 |  |  |
| Experienced both adulthood trauma and childhood trauma | 7 | 10 |  |  |
| Depression current |  |  | 0.32 | 1 |
| None | 30 | 35 |  |  |
| Yes | 8 | 4 |  |  |
| Depression lifetime |  |  | 0.84 | 1 |
| None | 30 | 29 |  |  |
| Yes | 8 | 10 |  |  |
| Persistent depressive disorder current |  |  | 0.49 | 1 |
| None | 31 | 35 |  |  |
| Yes | 7 | 4 |  |  |
| Persistent depressive disorder lifetime |  |  | 0.46 | 1 |
| None | 36 | 39 |  |  |
| Yes | 2 | 0 |  |  |
| PTSD Current |  |  | 0.43 | 1 |
| None | 19 | 24 |  |  |
| Yes | 19 | 15 |  |  |
| PTSD Lifetime |  |  | 0.99 | 1 |
| None | 1 | 0 |  |  |
| Yes | 37 | 39 |  |  |
| GAD Current |  |  | 0.23 | 1 |
| None | 35 | 39 |  |  |
| Yes | 3 | 0 |  |  |
| GAD lifetime |  |  | 0.91 | 1 |
| None | 38 | 39 |  |  |
| Yes | 0 | 0 |  |  |
| SAD Current |  |  | 1 | 1 |
| None | 35 | 36 |  |  |
| Yes | 3 | 3 |  |  |
| SAD Lifetime |  |  | 0.91 | 1 |
| None | 38 | 39 |  |  |
| Yes | 0 | 0 |  |  |
| Panic disorder current |  |  | 0.65 | 1 |
| None | 34 | 37 |  |  |
| Yes | 4 | 2 |  |  |
| Panic disorder lifetime |  |  | 0.91 | 1 |
| None | 38 | 39 |  |  |
| Yes | 0 | 0 |  |  |
| OCD current |  |  | 0.99 | 1 |
| None | 37 | 39 |  |  |
| Yes | 1 | 0 |  |  |
| OCD lifetime |  |  | 1 | 1 |
| None | 38 | 38 |  |  |
| Yes | 0 | 1 |  |  |
| SUD Current |  |  | 1 | 1 |
| None | 38 | 39 |  |  |
| Yes | 0 | 0 |  |  |
| SUD Lifetime |  |  | 0.41 | 1 |
| None | 33 | 37 |  |  |
| Yes | 5 | 2 |  |  |
| ADHD Current |  |  | 0.59 | 1 |
| None | 35 | 38 |  |  |
| Yes | 3 | 1 |  |  |
| ADHD Lifetime |  |  | 1 | 1 |
| None | 38 | 39 |  |  |
| Yes | 0 | 0 |  |  |
| GAD total | 7.22 | 5.49 | 0.23 | 0.39 |
| PCL total | 27.75 | 22.54 | 0.28 | 0.29 |
| MASQ total | 139.58 | 127.1 | 0.23 | 0.39 |
| SDS total | 12.67 | 8.85 | 0.10 | 0.39 |
| TEPS consummatory total | 32.08 | 32.56 | 0.8 | 0.804 |
| SHAPS FrankenTOT | 25.36 | 22.38 | 0.10 | 0.39 |
| SHAPS SnaithTOT | 2.67 | 43.44 | 0.11 | 0.39 |

GAD-7, Generalized anxiety disorder-7; PCL, PTSD Checklist for DSM-5; MASQ, Mood and Anxiety Symptom Questionnaire; SDS, Zung Self-Rating Depression Scale; TEPS consummatory, The Temporal Experience of Pleasure Scale-consummatory; SHAPS, Snaith-Hamilton Pleasure Scale
